# Integrated transcriptomic and metabolomic profiling reveals distinct molecular programmes across clinical subgroups of Sjögren’s disease and signatures associated with lymphoma development

**DOI:** 10.64898/2026.11.17.767332

**Authors:** Cristian Iperi, Shao Thing Teoh, Asimina Kakale, Silja Malkewitz, Marco Sprecher, Jacques-Eric Gottenberg, Yann Nguyen, Gaëtane Nocturne, Xavier Mariette, Bojana Müller-Durovic

**Affiliations:** Center for Experimental Rheumatology, University of Zurich and University Hospital of Zurich, Department of Rheumatology, Zurich-Schlieren, Switzerland; 2Rheumatology Department, EA 3432, Hôpitaux Universitaires de Strasbourg, Université de Strasbourg, France; Department of Rheumatology, Hôpital Bicêtre, Assistance Publique - Hôpitaux de Paris, Université Paris-Saclay, Paris, France; Université Paris-Saclay, Inserm, CEA; Immune diseases, microbiology and innovative therapies (IDMIT/UMRS1184); Fontenay-aux-Roses & Le Kremlin-Bicêtre, France

## Abstract

**Objectives:** Sjögren’s disease (SjD) is clinically heterogeneous and carries increased B-cell lymphoma risk. We aimed to identify immune-metabolic programmes underlying this heterogeneity and signatures associated with lymphoma.

**Methods:** We integrated whole-blood transcriptomics and plasma metabolomics from 344 ASSESS patients (306 matched profiles). Unsupervised (MOFA) and supervised (DIABLO) analyses characterised molecular programmes across three clinical clusters (BALS, HSA, LSAHS) and lymphoma status. Lymphoma-associated programmes were assessed by excluding lymphoma cases, modelling prevalent cases separately, and cluster-adjusted pathway scores. A transcriptomic score was developed using class-weighted LASSO regression and leave-one-out cross-validation.

**Results:** MOFA identified 18 latent factors grouped into five biological and one pharmacological theme, encompassing lymphocyte activation, interferon/redox stress, microbiota–tryptophan crosstalk and innate immunity. DIABLO linked BALS (IFN-high) to altered tryptophan metabolism and reduced microbiome-associated indoles; HSA (systemic inflammatory) to glutathione/redox metabolism, platelet activation and IL-18 signalling; and LSAHS to microbiome-associated metabolites and IL-10/AhR-related features. Lymphoma did not drive DIABLO factors; prevalent and incident cases showed higher cluster-adjusted IL-18/IL-1β and oxidative stress pathway scores, and prevalent cases an inverse IL-10/indole/KMO association. A transcriptomic score derived from 35 candidate genes, with eight retained by LASSO including IDH1 and genes involved in cellular-stress and adhesion, distinguished patients with prevalent or incident lymphoma from those without (AUC 0.796; permutation p=0.02). Incident lymphoma cases showed overlapping predicted probabilities before diagnosis.

**Conclusion:** Multi-omics integration identifies immune-metabolic programmes associated with SjD heterogeneity and lymphoma, highlighting inflammatory, redox and tryptophan-associated pathways. Detection of lymphoma-associated transcriptomic alterations before diagnosis supports prospective validation of blood-based biomarkers for risk stratification.

## Introduction

Sjögren’s disease (SjD) is a systemic autoimmune condition characterised by lymphocytic infiltration of exocrine glands, leading to sicca symptoms, fatigue, and a broad spectrum of extraglandular manifestations [1–3]. With a prevalence of approximately 0.1% in the general population and a strong female predominance, SjD represents one of the most common systemic autoimmune diseases [4–6]. Its clinical course is highly heterogeneous, ranging from predominantly glandular disease to severe systemic manifestations and lymphoma [5,7,8]. This heterogeneity has hampered the development of effective therapies and prognostic tools [9–12]. Although clinically distinct patient subgroups have been identified, the molecular programmes underlying these disease states and those associated with progression towards lymphoma remain poorly understood. SjD carries one of the highest lymphoma risks among autoimmune diseases, with an approximately 15-to 20-fold increase in B-cell non-Hodgkin lymphoma, predominantly mucosa-associated lymphoid tissue (MALT) lymphoma, which represents its most severe complication and occurs in approximately 5–10% of patients [13,14]. No validated biomarker panel exists to identify patients at highest risk prospectively, a major unmet clinical need.

Efforts to stratify SjD patients beyond clinical phenotype have increasingly relied on molecular profiling of peripheral blood. Transcriptomic studies have identified recurrent subgroups defined by IFN-I pathway activation, lymphocyte subset composition, and inflammatory gene expression programmes [8,15], paralleling similar stratification efforts in systemic lupus erythematosus and other systemic autoimmune diseases [16,17]. The ASSESS cohort, a prospective multicentre French cohort with a 20-year follow-up, has recently defined three clinical clusters termed B-cell active disease and low symptom burden (BALS), high systemic disease activity (HSA), and low systemic disease activity and high symptom burden (LSAHS), using clinical and serological variables [8]. A complementary study further characterised transcriptomic differences across these clusters in the ASSESS cohort [18]. However, transcriptomic profiling alone cannot provide a complete picture of the biological processes underlying SjD heterogeneity, while metabolic dysregulation is increasingly recognised in autoimmune diseases [19–22]. Key metabolic programmes, such as tryptophan catabolism along the kynurenine–AhR–IL-10 axis or glutathione-dependent redox regulation, are shaped by post-translational control, enzymatic activity and microbiome–host interactions, and therefore require direct measurement at the metabolite level.

The plasma metabolome therefore offers an orthogonal view of immune and metabolic states. Plasma metabolomics, now feasible at scale with untargeted liquid chromatography– high-resolution mass spectrometry (LC-HRMS), has not previously been integrated with transcriptomic data in the ASSESS cohort. Here, we address this gap by integrating plasma metabolomics with transcriptomics through unsupervised (MOFA) and supervised (DIABLO) multi-omics frameworks to ask which molecular programmes distinguish clinically meaningful SjD subgroups, and which of these programmes identify patients progressing towards lymphoma.

## Methods

Full methodological details are provided in the Supplementary Methods. In brief, we studied patients from the ASSESS cohort, a prospective multicentre French cohort (15 tertiary centres, recruitment 2006–2009, with 20-year follow-up) designed to identify predictors of systemic complications and lymphoma in Sjögren’s disease [8,18,23]. All patients fulfilled the 2002 AECG criteria and gave written informed consent; the study was approved by the Ethics Committee of Bichat Hospital (Approval dossier N° 0711468). Of 344 patients with plasma metabolomic data, 306 also had matched whole-blood transcriptomic data. Patients belonged to three previously defined clinical clusters: BALS (n=160), HSA (n=138) and LSAHS (n=46) [8]. Seventeen patients had lymphoma at enrolment (prevalent lymphoma, pLym) and 12 developed lymphoma during follow-up (incident lymphoma, iLym). Of these, 15 pLym and all 12 iLym had matched transcriptomic data and were included in DIABLO and transcriptome-based analyses.

Gene expression in whole blood was measured by microarray (GSE140161) [24], and the 5,000 most variable genes were retained. Plasma metabolites were measured by untargeted LC-HRMS in positive and negative ionisation modes, yielding 1,413 and 802 features, respectively. Quality control, normalisation, metabolite annotation against public spectral databases [25–30], and adjustment for technical batch effects and corticosteroid use with the limma package [31] are detailed in the Supplementary Methods.

We used two complementary multi-omics integration approaches (**Figure 1**). Multi-omics factor analysis (MOFA) is an unsupervised method that identifies latent factors capturing major axes of variation within and across the transcriptome, metabolome and clinical data, without knowledge of the clinical clusters, and was applied to all 344 patients [32]. DIABLO, a supervised method identifying combinations of genes and metabolites associated with the clinical clusters, was applied to the 306 patients with complete data [33]. The number of factors was selected using a composite score, evaluated for MOFA across five runs with different random seeds. To investigate the influence of lymphoma on the identified molecular programmes, DIABLO was run (i) on all patients, (ii) after excluding pLym patients, iLym patients or both, and (iii) with pLym patients assigned to a dedicated fourth group. Each scenario was compared with 200 refits using randomly selected, cluster-matched lymphoma-free patients, and feature overlap between the three- and four-group models was quantified. Factors were interpreted based on their strongest contributing features (scaled loadings >0.5 or <−0.5), pathway enrichment (EnrichR) and metabolite annotations [34], as detailed in the Supplementary Methods. Programmes highlighted by DIABLO were then tested independently with whole-transcriptome pathway scores (Gene Ontology gene sets and those used to annotate the DIABLO factors) and selected genes and metabolites, using linear models adjusted for clinical cluster with Benjamini–Hochberg correction.

**Figure 1.**
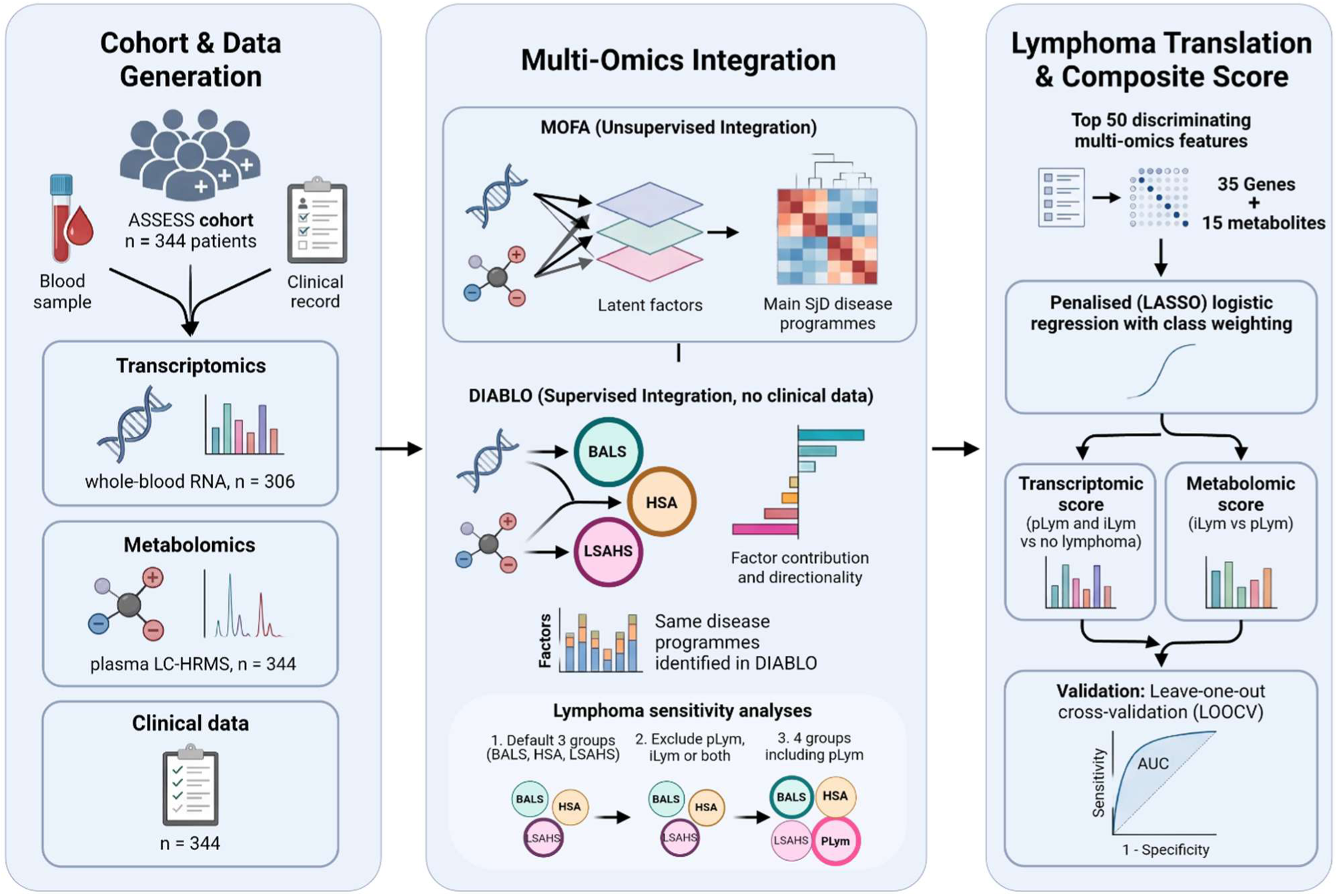
Study design and multi-omics analytical workflow. *Left panel:* Cohort and data generation. Whole-blood and plasma samples and clinical data were collected from 344 patients of the ASSESS cohort, generating whole-blood transcriptomic profiles (Affymetrix microarrays; n=306), plasma metabolomic profiles (LC-HRMS, positive and negative ionisation modes; n=344), and clinical data (n=344). *Centre panel:* Multi-omics integration. Unsupervised multi-omics factor analysis (MOFA) was applied across all three data types, identifying latent factors capturing major axes of molecular and clinical variation. Supervised integration (DIABLO) was applied independently to the 306 patients with matched transcriptomic and metabolomic profiles, without integration of clinical data, to identify molecular programmes associated with three previously defined patient clusters (Nguyen et al. [8]): BALS (B-cell active disease and low symptom burden), HSA (high systemic disease activity), and LSAHS (low systemic disease activity and high symptom burden). The factors identified by DIABLO recapitulated the disease programmes identified by MOFA. To assess the influence of lymphoma, DIABLO was run with the three clinical clusters, after excluding patients with prevalent lymphoma (pLym; diagnosed at cohort recruitment), incident lymphoma (iLym, diagnosed during follow-up) or both, each compared with 200 random exclusions of cluster-matched lymphoma-free patients, and with pLym as a dedicated fourth group. Programmes highlighted by DIABLO were then tested with cluster-adjusted whole-transcriptome pathway scores. *Right panel:* Lymphoma-associated score development. Penalised class-weighted LASSO logistic regression was applied to the top 50 discriminating multi-omics features to derive two scores. The transcriptomic score was designed to distinguish patients with lymphoma, either prevalent (pLym) or incident (iLym), from those without lymphoma, while the metabolomic score distinguished iLym from pLym. Performance was assessed by leave-one-out cross-validation (LOOCV) using the area under the ROC curve (AUC).

Genes and metabolites differing across five mutually exclusive groups (lymphoma-free BALS, HSA and LSAHS, pLym and iLym) were screened using Kruskal–Wallis tests. The 50 most discriminating features were used to build a transcriptomic score distinguishing patients with lymphoma from those without (pLym and iLym vs lymphoma-free; 35 candidate genes) and a metabolomic score distinguishing incident from prevalent lymphoma (iLym vs pLym; 15 candidate features) using penalised (LASSO) logistic regression with class weighting. Performance was assessed by leave-one-out cross-validation (LOOCV), with the area under the ROC curve (AUC), bootstrap 95% confidence intervals, and a 1,000-permutation test against chance.

Analyses were performed in R v4.4.3. Group comparisons used Kruskal–Wallis, Dunn (Benjamini–Hochberg-adjusted) and Wilcoxon tests; two-sided p-values <0.05 were considered significant.

## Results

### Integrated multi-omics profiling identifies distinct immune and metabolic programmes in Sjögren’s disease

To identify the major molecular programmes underlying SjD, we integrated whole-blood transcriptomics (n=306), positive and negative ionisation-mode plasma metabolomics (n=344), and clinical data from 344 ASSESS patients. The study design and analytical workflow are summarised in **Figure 1**. MOFA is an unsupervised approach that identifies latent factors, representing patterns of genes, metabolites and clinical variables that vary together across patients, without prior knowledge of their clinical subgroup. The number of factors was selected based on model stability across independent runs, and their biological interpretation was based on the most strongly contributing features and pathway enrichment. MOFA identified 18 latent factors collectively explaining 67.8%, 39.9%, 16.1%, and 2.1% of the variance within the transcriptomic, positive metabolomic, negative metabolomic, and clinical data layers, respectively. The transcriptomic layer was the dominant contributor, with Factor 1 alone explaining 23.6% of transcriptomic variance, while the metabolomic layers contributed distinct axes of molecular variation, predominantly in Factors 2–8 (**Figure 2A-B**). In contrast to the transcriptomic and metabolomic layers, the clinical layer showed substantially lower variance explained across the 18 factors (2.1%), with Factor 6 representing the primary axis of clinical co-variation.

**Figure 2.**
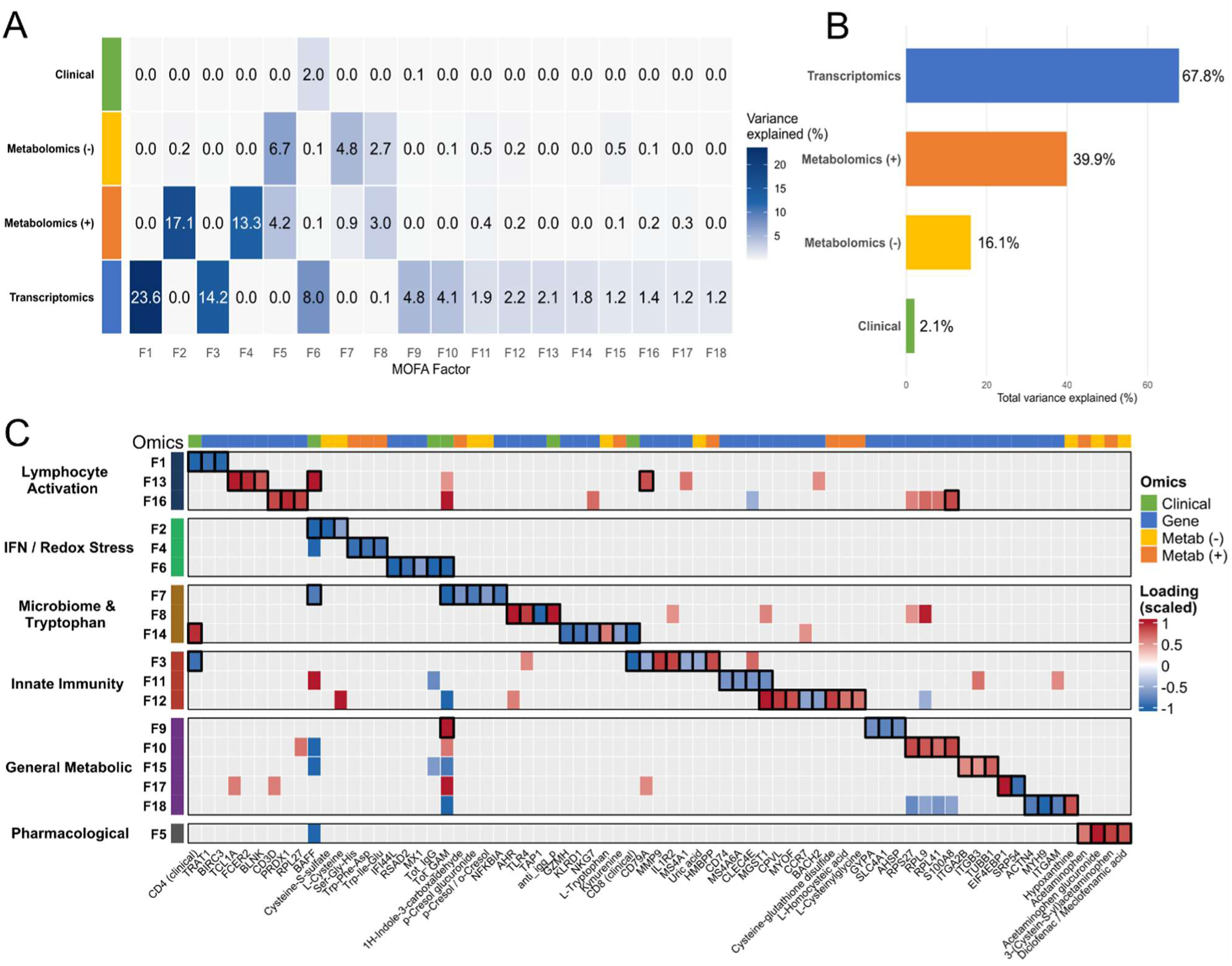
MOFA identifies 18 latent factors representing distinct immune and metabolic programmes in Sjögren’s disease. **A)** Heatmap of the percentage of variance explained by each of the 18 MOFA factors within each omic layer (Clinical, Metabolomics (−), Metabolomics (+), Transcriptomics). Colour intensity and the value in each cell indicate the percentage of variance explained by that factor within that specific layer. **B)** Total variance explained (%) within each omic layer, summed across all 18 factors, illustrating the relative contribution of each data modality to the overall multi-omics model. **C)** Heatmap of the top contributing features supporting the biological annotation of each MOFA factor. Rows represent the 18 factors, grouped into five biological themes (Lymphocyte Activation, IFN/Redox Stress, Microbiome & Tryptophan, Innate Immunity, General Metabolic) and one pharmacological theme, indicated by the coloured side bar. Columns represent selected genes, metabolites, and clinical variables, coloured by their omic layer of origin (Omics annotation bar: Clinical, Gene, Metabolomics (−), Metabolomics (+)). Cell colour indicates the loading score for each feature on each factor, scaled to the maximum absolute loading across all views within that factor (range −1 to +1; blue = negative, red = positive); grey cells indicate that the feature was not among the top loadings for that factor. Black borders mark the factor(s) for which a feature was selected as a top contributing loading used to support the biological annotation.

Based on top-loading features and pathway enrichment (EnrichR), the 18 MOFA factors were grouped into five biological themes and one pharmacological theme (**Figure 2C**). A lymphocyte activation programme encompassed CD4+ T-cell and B-cell activation-related factors. Closely related was an IFN-I/redox stress programme, combining canonical interferon signalling with a recurring sulfur-redox/glutathione theme across multiple independent factors, highlighting metabolic features not captured by transcriptomics alone. A third, microbiota–tryptophan programme linked natural killer cell (NK) and CD8+ T cell (NK/CD8) cytotoxicity, indoleamine 2,3-dioxygenase (IDO)-mediated tryptophan catabolism, and indole-associated metabolites potentially related to AhR signalling. Innate immunity emerged as a fourth programme, capturing the neutrophil-to-lymphocyte ratio, consistent with previously described transcriptomic stratification in SjD [15], alongside myeloid stress and respiratory burst regulation. Finally, general metabolic factors reflected ribosomal translation, erythroid biology, and platelet activation, while a separate factor captured pharmacological signatures of anti-inflammatory treatments. Collectively, these 18 factors define the principal molecular axes identified in Sjögren’s disease, spanning adaptive and innate immune activation, redox stress, microbiota-immune crosstalk, and metabolic reprogramming (**Figure 2C**; **Supplementary Table 1**).

### Distinct molecular programmes characterise the three clinical Sjögren’s disease subgroups

To determine how the molecular programmes identified by unsupervised analysis relate to clinically defined Sjögren’s disease phenotypes, we applied DIABLO to discriminate between BALS, HSA, and LSAHS. DIABLO identified 16 multi-omics factors that broadly recapitulated the biological programmes identified by MOFA, including IFN-I/tryptophan metabolism, cytotoxic NK/CD8 activity, myeloid activation, and glutathione/redox metabolism, while revealing their differential associations with the three clinical clusters (**Figure 3A-C**).

**Figure 3.**
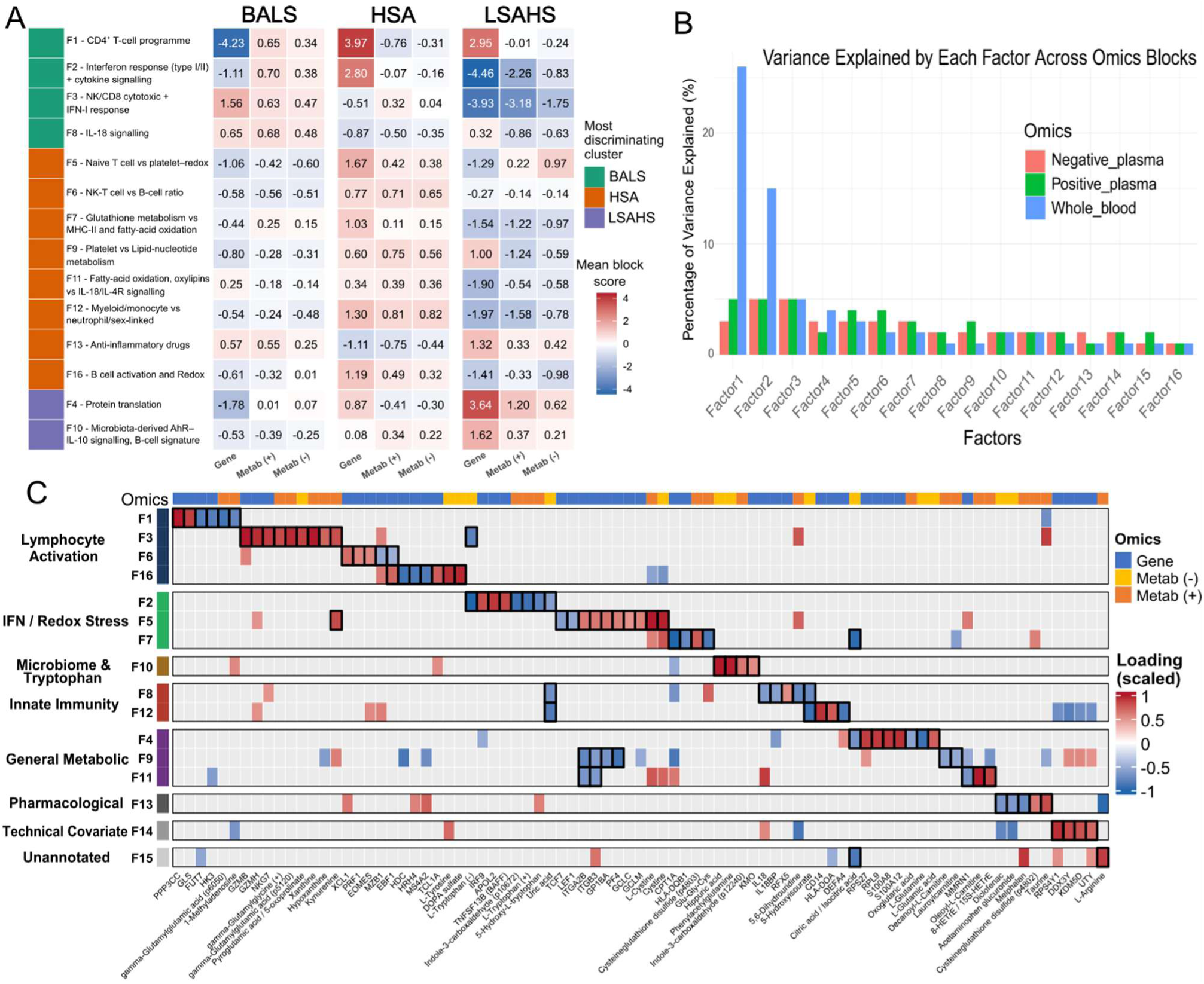
DIABLO identifies molecular programmes associated with the three clinical Sjögren’s disease subgroups. **A)** Heatmap of native, per-block mean latent scores for the selected DIABLO factors across the three clinical clusters (BALS, HSA and LSAHS) and the three omic blocks (Transcriptomics, Metabolomics (+), Metabolomics (−)). The unannotated and technical factors were excluded from the heatmap. Rows are grouped according to the clinical cluster with which each factor shows the strongest association from the heatmap, indicated by the coloured side bar; cell colour indicates the magnitude and direction of the mean latent scores (blue = negative, red = positive) for that block within that factor. **B)** Bar plot showing the percentage of variance explained by each of the 16 DIABLO factors within each omic block (Transcriptomics, Metabolomics (+), Metabolomics (−)), illustrating the variation captured by each factor within the individual data modality. **C)** Heatmap of the top contributing features selected to support the biological annotation of each DIABLO factor, presented as in Figure 2C. Rows represent the 16 factors, grouped into biological themes as indicated by the coloured side bar. Columns represent selected genes and metabolites, coloured by their omic layer of origin. Cell colour indicates the loading score for each feature on each factor, scaled to the maximum absolute loading across all views within that factor (range −1 to +1; blue = negative, red = positive); grey cells indicate the feature was not among the top loadings for that factor. Black borders mark the factor(s) for which a feature was selected as a top contributing loading used to support the biological annotation.

BALS was characterised by prominent cytotoxic NK/CD8 and tryptophan-associated signatures, alongside the previously described IFN-I-high phenotype. Factor 3 captured an IFN-I-associated programme with combined NK/CD8 activity, kynurenine-associated metabolites, and purine catabolism. Together, these findings link the BALS phenotype to perturbation of tryptophan metabolism, consistent with the IFN-high subgroups previously described in Sjögren’s disease [15] and other systemic autoimmune diseases [20]. BALS also showed an inverse association with Factor 10, characterised by KMO expression, IL-10-related features, and microbiome-associated metabolites, including hippuric acid, phenylacetylglutamine and indole-3-carboxaldehyde. This opposing pattern suggests distinct associations of kynurenine-and indole-related tryptophan metabolism in the BALS phenotype. BALS was also negatively associated with Factor 4, representing a translational/ribosomal programme, consistent with previously reported alterations in ribosomal gene expression in SjD B cells [35]. Factor 8, capturing IL-18-related features, was also positively associated with BALS, suggesting an additional innate immune component alongside the interferon axis.

HSA displayed a distinct signature of systemic inflammatory activation. Notably, Factor 2 showed different contributions from the two omic layers. Its transcriptomic component, comprising type I and type II IFN and cytokine signalling, was more pronounced in HSA than in BALS, suggesting that IFN-I-associated transcriptional features are not restricted to BALS but are embedded in a broader inflammatory signature in HSA. In contrast, its metabolomic component, reflecting tryptophan-associated programmes, was more strongly associated with BALS, similar to the IFN-I and NK/CD8 signature of Factor 3. In HSA, Factor 7 linked glutathione metabolism with MHC class II antigen presentation and fatty acid oxidation. Platelet-associated features emerged as a prominent theme, captured independently by Factors 5 and 11, the latter also including IL-18/IL-4R-related features. A myeloid programme (Factor 12) further distinguished HSA, with positive scores associated with monocyte markers including CD14. Finally, a treatment-associated factor (Factor 13) also differed between the clinical clusters.

LSAHS was characterised by a positive association with Factor 10, encompassing microbiome-associated metabolites and features related to the AhR/IL-10 axis. LSAHS also showed the strongest positive association with Factor 4, representing translational/ribosomal activity, whereas this association was weaker in HSA and negative in BALS at the transcriptomic level. The factor loadings for all 16 DIABLO factors are provided in **Supplementary Table 2.**

### Lymphoma patients share the multi-omic architecture of their clinical cluster but display an IL-18/IL-1β and oxidative stress transcriptional programme

Patients with lymphoma were unevenly distributed across clinical clusters, with most lymphoma cases, particularly prevalent ones, classified as HSA (**Table 1**). We therefore asked whether lymphoma patients share the molecular profile of their cluster or display additional lymphoma-associated features. We first assessed whether lymphoma patients shaped the DIABLO factor structure by refitting the model after excluding patients with prevalent lymphoma (pLym, n=15), patients who developed lymphoma during follow-up (incident lymphoma, iLym, n=12), or both (n=27). Each exclusion was compared with 200 random exclusions of matched lymphoma-free patients. For all 16 factors and all three exclusion scenarios, similarity to the original model remained within the range expected from random exclusion (all FDR ≥ 0.35; **Supplementary Figure 1**). Thus, the multi-omic architecture of the clinical clusters was not driven by patients with lymphoma.

**Table 1.** Clinical characteristics of patients with prevalent and incident lymphoma in the ASSESS cohort.

|  |  | **Prevalent lymphoma (n=17)** | **Incident lymphoma (n=12)** |
|---|---|---|---|
| Age, years | Mean ± SD | 63.06 ± 10.15 | 58.75 ± 5.93 |
| Female sex | n (%) | 16 (94.1) | 11 (91.7) |
| Time from sampling to lymphoma diagnosis, years* | Mean ± SD | – | 9.53 ± 6.28 |
| **Clinical cluster** |  |  |  |
| BALS | n (%) | 2 (11.8) | 5 (41.7) |
| HSA | n (%) | 15 (88.2) | 7 (58.3) |
| LSAHS | n (%) | – | – |
| **Lymphoma subtype** |  |  |  |
| MALT lymphoma | n (%) | 8 (47.1) | 6 (50.0) |
| Marginal zone lymphoma (other) | n (%) | 4 (23.5) | – |
| DLBCL | n (%) | 4 (23.5) | 4 (33.3) |
| Small B-cell lymphoproliferative disorder | n (%) | – | 2 (16.7) |
| EBV-associated B-cell lymphoma | n (%) | 1 (5.9) | – |
| **Treatment** |  |  |  |
| Rituximab monotherapy | n (%) | – | 3 (25.0) |
| Rituximab + chemotherapy | n (%) | 1 (5.9) | 5 (41.7) |
| Rituximab + belimumab | n (%) | 1 (5.9) | – |
| Chemotherapy, unspecified | n (%) | – | 1 (8.3) |
| Targeted therapy | n (%) | – | 2 (16.7) |
| Immunoglobulin replacement | n (%) | 1 (5.9) | – |
| NA | n (%) | 14 (82.4) | 2 (16.7) |

n: Number of patients. \*For two patients only the year of lymphoma diagnosis was available; the date was set to 1 July of that year. Of the 17 patients with prevalent lymphoma, 15 had transcriptomic data.
DLBCL, diffuse large B-cell lymphoma; EBV, Epstein–Barr virus; MALT, mucosa-associated lymphoid tissue; NA, not available.

Assigning patients with prevalent lymphoma to a dedicated fourth cluster (pLym) largely preserved the DIABLO factor structure, with a median feature overlap of 76.1% between the original three-class and four-class models (**Figure 4A-B**), comparable to that observed when the same number of randomly selected patients was assigned to a fourth class (**Supplementary Figure 1**). Within this model, pLym patients showed stronger associations with several molecular programmes already observed in HSA, while also displaying additional lymphoma-associated features. The IFN/cytokine-associated programme (Factor 2) and nucleotide catabolism-associated programme (Factor 1) showed their strongest associations with pLym. Major transcriptomic contributors to Factor 1 included *FUT7*, previously implicated in lymphoma development [36]. The platelet and glutathione-associated programme (Factor 5) was also more strongly associated with pLym than with HSA. Conversely, Factor 10, capturing the IL-10/indole/KMO axis, showed its strongest negative association with pLym. Factor 15 further distinguished the lymphoma-associated profile, with bile acid-related metabolites associated with pLym, in contrast to the mast cell-associated markers in BALS. Notably, not all redox-associated programmes showed a stronger association with pLym. The glutathione metabolism factor (Factor 7) remained more strongly associated with HSA than with pLym (**Supplementary Table 3**). Finally, Factor 8 showed a strong association with pLym. This factor was characterised by *INPP4B* and *IER3* as top transcriptomic contributors, alongside *IL18*, *IL18BP* and *IL1B*, together with metabolomic features associated with nitrosative stress, including 3-nitrotyrosine and uric acid.

**Figure 4.**
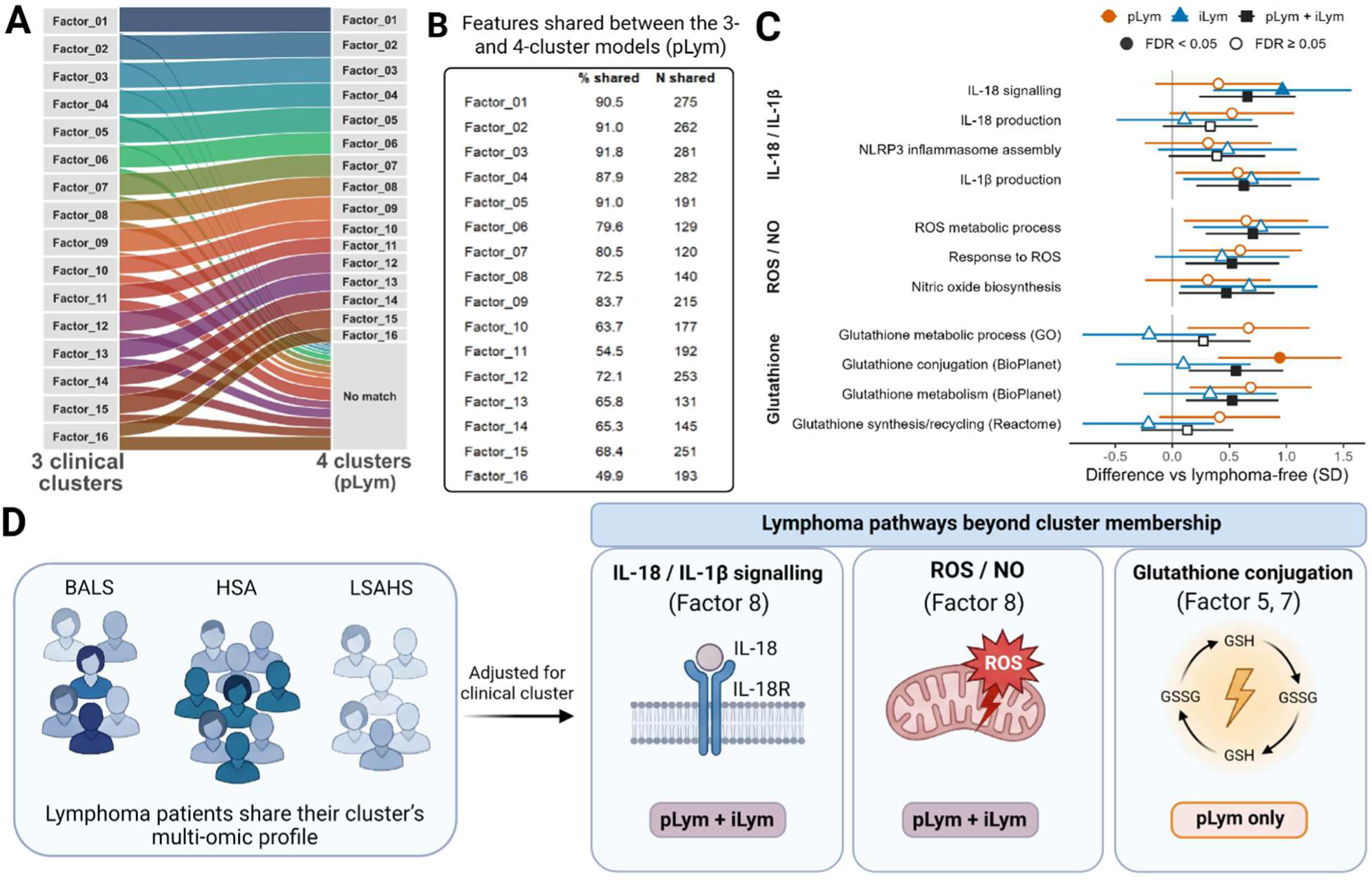
Influence of lymphoma on DIABLO-derived molecular programmes and lymphoma-associated pathways. **A)** Alluvial plot comparing factor composition between the original three-class DIABLO model based on the clinical clusters (left) and the four-class model, in which patients with prevalent lymphoma were assigned to a dedicated fourth cluster (pLym; right). Ribbons connect corresponding factors, with width representing the overlap of top-contributing transcripts and metabolites; unmatched features are shown as ‘no match’. **B)** Number and percentage of top-contributing features of each factor of the three-class model that were shared with the four-class model. **C)** Whole-blood pathway scores in patients with prevalent (pLym) and incident (iLym) lymphoma, and in both groups combined (pLym + iLym), compared with lymphoma-free patients. Points show the difference in pathway score, adjusted for clinical cluster and expressed in standard deviations (SD) of lymphoma-free patients, with 95% confidence intervals; filled symbols indicate a false discovery rate (FDR) < 0.05. Gene sets were obtained from Gene Ontology (GO), BioPlanet 2019 and Reactome 2024. **D)** Schematic summary. Patients with lymphoma (darker icons) are distributed within the clinical clusters and share their multi-omic profile, while displaying IL-18/IL-1β and reactive oxygen species/nitric oxide (ROS/NO) transcriptional programmes in both prevalent and incident lymphoma, and increased glutathione conjugation-related transcription in prevalent lymphoma only. Factor numbers indicate the DIABLO factors from which each programme was derived.

Because factor associations in a supervised model can overstate differences in small groups, we tested two pre-specified programmes independently of DIABLO: IL-18/IL-1β signalling (Factor 8), previously linked to SjD and SjD-associated lymphoma [37,38], and glutathione metabolism (Factors 5 and 7)[39], a redox theme recurring across MOFA and DIABLO factors. As iLym patients were not labelled in DIABLO, they provided an independent test. Using the whole transcriptome, we scored gene sets related to IL-18/IL- 1β and inflammasome activity, glutathione metabolism, including those originally used to annotate Factors 5 and 7, and reactive oxygen and nitrogen species, and compared patients with prevalent, incident or any lymphoma with lymphoma-free patients after adjustment for clinical cluster (**Figure 4C; Supplementary Table 4**). Patients with lymphoma showed higher IL-18 signalling, IL-1β production and reactive oxygen and nitric oxide-related scores. These differences were similar in prevalent and incident cases and were not explained by cluster membership, as these pathway scores did not differ between HSA and BALS. Consistently, lymphoma patients showed reduced expression of *IL18BP*, which encodes the endogenous IL-18 inhibitor, and of *INPP4B*. In contrast, glutathione-related transcription, particularly glutathione conjugation, was increased in prevalent but not in incident lymphoma. At the metabolomic level, the redox-and nitrosative stress-related metabolites contributing to these programmes did not differ according to lymphoma status (**Supplementary Table 5**). Together, these findings indicate that, beyond the immune-metabolic profile shared with their clinical cluster, patients with lymphoma display an IL-18/IL-1β and oxidative stress transcriptional programme that is already detectable before lymphoma diagnosis (**Figure 4D**).

### A compact transcriptomic score distinguishes patients with prevalent or incident lymphoma from those without lymphoma

Since this transcriptional programme was shared by patients with prevalent and incident lymphoma, we next asked whether a limited set of features, selected without any a priori biological criterion, could distinguish patients according to lymphoma status. Features were ranked by Kruskal–Wallis screening across five mutually exclusive groups (lymphoma-free BALS, HSA and LSAHS, pLym and iLym), and the top 50 (35 genes and 15 metabolites) were used to develop separate transcriptomic and metabolomic scores.

A composite transcriptomic score (Score 1) integrating 35 candidate genes was designed to distinguish patients with prevalent or incident lymphoma (pLym and iLym) from patients without lymphoma (BALS, HSA, LSAHS). Score 1 achieved an AUC of 0.796 (95% CI 0.700–0.887), significantly exceeding chance performance (permutation test p=0.02; 1,000 permutations). Notably, patients who subsequently developed lymphoma showed elevated LOOCV-derived predicted probabilities that overlapped with those observed in patients with prevalent lymphoma (**Figure 5A-C**). As all iLym samples were collected before diagnosis, these transcriptomic features were detectable before clinical lymphoma. LASSO feature selection identified eight genes as the primary discriminators (**Figure 5D**): *IDH1, WDR41* (positive coefficient, higher in pLym + iLym), and *DSC1*, DPP4, *PATJ, ITGA6, USP44 and ZNF860* (all negative coefficients, lower in pLym + iLym).

**Figure 5.**
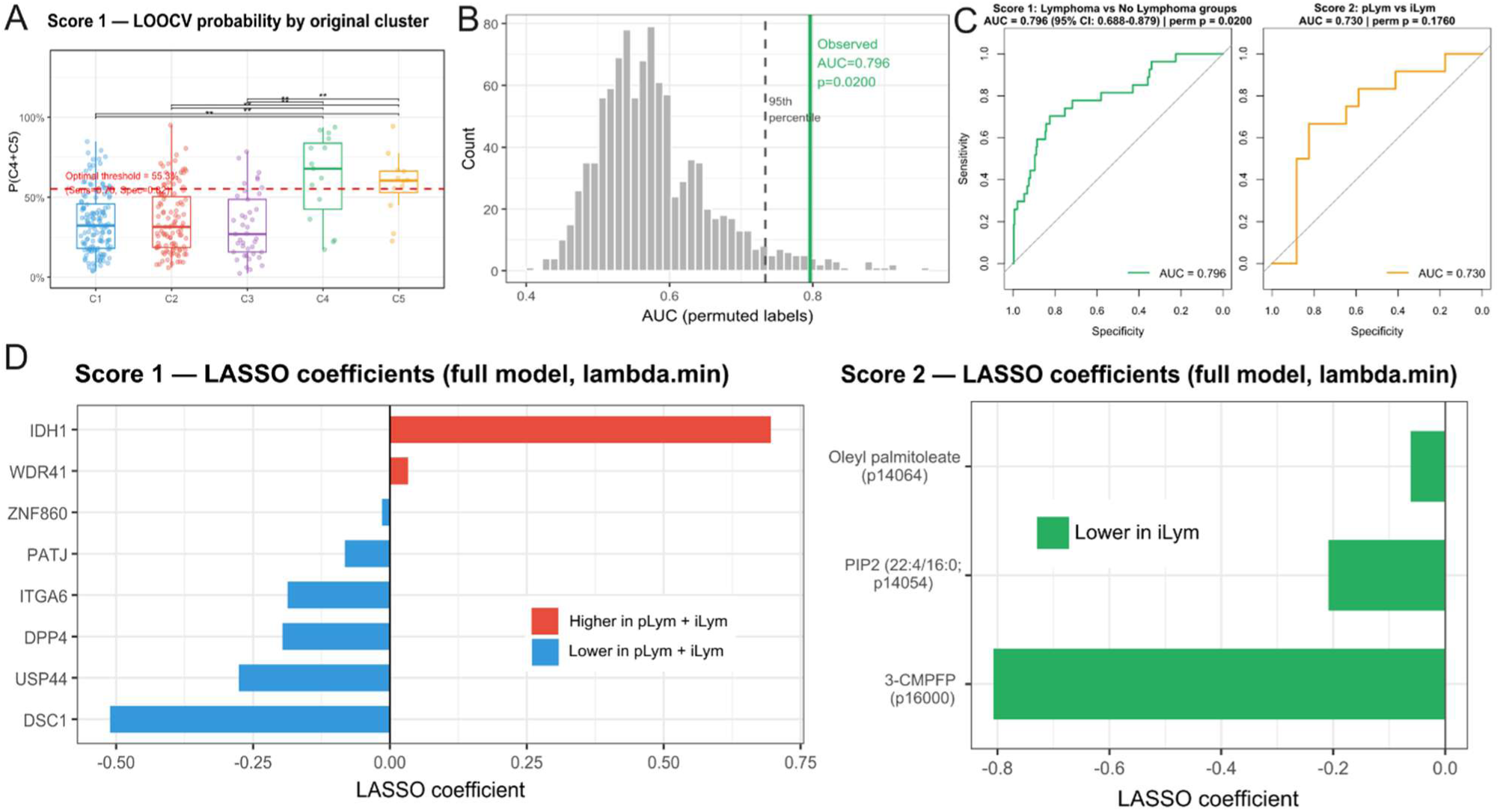
Transcriptomic and metabolomic scores associated with prevalent and incident lymphoma. **A)** LOOCV-derived predicted probabilities of lymphoma for transcriptomic Score 1 across the original clinical groups (BALS, HSA, LSAHS, pLym, iLym), illustrating the position of incident lymphoma patients (iLym) relative to prevalent lymphoma patients (pLym) and non-lymphoma clusters. The dashed line indicates the 95^th^ percentile of the AUC distributions with permuted labels. **B)** Null distribution of AUC values from 1,000 permutations for Score 1, with the observed AUC and 95^th^ percentile indicated. **C)** ROC curves for Score 1 (prevalent and incident lymphoma versus no lymphoma) and Score 2 (incident versus prevalent lymphoma), with their respective AUC values and permutation p-values. The 95% bootstrap confidence interval is additionally shown for Score 1. **D)** LASSO regression coefficients for the genes retained in the final Score 1 model (left) and metabolites retained in the final Score 2 model (right). Positive coefficients indicate an association with higher odds of belonging to the lymphoma group (pLym + iLym versus no lymphoma) for Score 1, whereas negative coefficients indicate lower odds. For Score 2, negative coefficients indicate an association with lower odds of incident lymphoma (iLym) relative to prevalent lymphoma (pLym).

A complementary plasma metabolomic score (Score 2), developed from 15 candidate features, was used to distinguish patients with prevalent from those with incident lymphoma. Score 2 achieved an AUC of 0.730 (95% CI 0.520–0.907), but did not significantly exceed chance (permutation test p=0.196; 1,000 permutations). LASSO feature selection retained three metabolomic features in the final model (**Figure 5D**; MS1 annotation): PIP2(22:4/16:0), oleyl palmitoleate and 3-carboxy-4-methyl-5-pentyl-2-furanpropanoic acid (3-CMPFP). All three had negative coefficients, the largest being for 3-CMPFP. Given the limited sample size, no conclusion can be drawn on its discriminatory utility.

Together, these results identify a transcriptomic signature associated with both prevalent and incident lymphoma, detectable before clinical diagnosis in patients who subsequently developed lymphoma. The metabolomic analysis additionally suggests differences between prevalent and incident lymphoma, although these findings require validation in larger cohorts.

## Discussion

In this study, we applied integrated whole-blood transcriptomic and plasma metabolomic profiling to the ASSESS cohort to define the major molecular programmes underlying disease variation and lymphoma status. MOFA and DIABLO converged on five biological programmes and one pharmacological programme, supporting the consistency of these axes across analytical approaches. BALS was characterised by IFN-I activation and altered tryptophan metabolism, HSA by glutathione/redox metabolism, platelet activation and IL-18/IL-4R signalling, and LSAHS by microbiome-associated metabolites and IL-10/AhR-related features. Patients with lymphoma shared the multi-omic architecture of their clinical cluster but displayed an IL-18/IL-1β and oxidative stress transcriptional programme, and transcriptomic features shared by prevalent and incident cases were detectable before clinical diagnosis.

The dominant transcriptomic signature of BALS was a strong IFN-I response, consistent with the IFN-high signature previously described in peripheral blood from SjD patients [8,15] and other systemic autoimmune diseases [16,20]. Our integrated analysis further linked the BALS molecular profile to lower circulating levels of selected indole metabolites, which can originate from microbial tryptophan metabolism and can engage the aryl hydrocarbon receptor (AhR). AhR activation by indole derivatives has been implicated in immunoregulatory pathways, including IL-10 production by regulatory B cells and Tr1 cells [40]. Thus, inflammatory activation in BALS may be accompanied by alterations in the microbiota–tryptophan–AhR–IL-10 immunoregulatory axis, potentially limiting compensatory anti-inflammatory mechanisms [41]. Furthermore, lower tryptophan and higher kynurenine levels in BALS (Factor 3), consistent with IFN-induced IDO1-mediated tryptophan catabolism, may also reduce the availability of tryptophan for microbial indole production [42]. These findings highlight the AhR–IL-10 axis as a candidate for future mechanistic and therapeutic investigation.

HSA displayed a distinct inflammatory and metabolic profile dominated by glutathione/redox metabolism, platelet activation and IL-18/IL-4R signalling, together with myeloid activation and antigen-presentation programmes. Factor 7 linked glutathione-related metabolites with MHC-II antigen presentation and fatty acid oxidation, suggesting altered glutathione turnover and redox regulation in the context of systemic immune activation. Platelet activation emerged independently across two DIABLO factors (Factors 5 and 11), consistent with growing evidence that platelets contribute to systemic autoimmunity through release of inflammatory mediators and interaction with immune cells [43]. A further myeloid programme including CD14 and IL-18/IL-4R signalling reinforced the association of HSA with innate immune activation. Notably, the type I and type II IFN and cytokine transcriptional component of Factor 2 was more pronounced in HSA than in BALS, whereas its tryptophan-associated metabolomic component and the cytotoxic NK/CD8 programme of Factor 3 were more characteristic of BALS. This suggests that IFN activation is not restricted to BALS but, in HSA, is embedded in a broader inflammatory response involving IL-18 signalling, myeloid activation and antigen presentation. These molecular programmes, together with treatment-associated differences, may reflect the higher systemic disease activity and treatment burden characterising this subgroup.

Removal of patients with prevalent lymphoma, incident lymphoma or both did not alter the DIABLO factors beyond what was expected from removing the same number of lymphoma-free patients, indicating that the multi-omic architecture of the clinical clusters is not driven by patients with lymphoma. The preserved factor composition in the four-class model further suggests that the lymphoma-associated profile emerges mainly within this architecture.

Beyond this shared architecture, the IL-18/IL-1β and oxidative stress transcriptional programme highlighted by Factor 8 is of particular interest, as IL-18 has been implicated in lymphoma biology and SjD-associated inflammation, including inflammasome activation [37,38]. Among the genes most associated with this factor, *INPP4B* encodes a phosphatase that counteracts PI3K/AKT signalling, and its downregulation in patients with lymphoma is consistent with observations in B-cell lymphoma [44]. Pathway analyses independent of DIABLO confirmed this programme in both prevalent and incident lymphoma, together with reduced expression of *IL18BP*, the endogenous IL-18 inhibitor. As these scores did not differ between HSA and BALS, the programme appears to reflect lymphoma status rather than cluster membership, and may contribute to the molecular environment accompanying lymphoma development in SjD. In contrast, redox-and nitrosative stress-related plasma metabolites did not differ according to lymphoma status, indicating that this programme is detectable at the transcriptional but not at the circulating metabolite level. Glutathione conjugation-related transcription was increased in prevalent but not in incident lymphoma, suggesting that this feature may accompany established disease rather than precede it.

A particularly relevant finding was the similarity of transcriptomic signatures between patients with prevalent lymphoma and those who subsequently developed lymphoma, whereas their metabolomic profiles showed greater divergence. Consistently, IL-18/IL-1β and oxidative stress pathway scores were similar in prevalent and incident cases. Importantly, all incident lymphoma patients were sampled before lymphoma diagnosis and initiation of lymphoma-directed treatment, supporting the presence of lymphoma-associated transcriptional alterations before clinical transformation [18]. This suggests that circulating transcriptomic profiles may capture early molecular changes associated with lymphomagenesis, whereas metabolomic signatures may additionally reflect the systemic metabolic state associated with established lymphoma.

Interestingly, Factor 10, comprising microbiome-associated metabolites and IL-10/AhR- related features, was positively associated with LSAHS but negatively associated with BALS, most strongly in patients with prevalent lymphoma. Notably, LSAHS was the only clinical subgroup in which no lymphoma cases developed during follow-up. Given the role of microbiota-derived indoles in AhR-dependent IL-10 production, whether this programme protects against lymphoma development remains to be investigated.

The eight genes retained in the transcriptomic score mapped onto several biological themes. Altered central carbon and redox metabolism was reflected by *IDH1*, which encodes the cytosolic NADP+-dependent isocitrate dehydrogenase and has been implicated in haematological malignancies [45]. *IDH1*, which was higher in patients with lymphoma, also contributes to NADPH production and redox homeostasis, but its role in SjD-associated lymphomagenesis remains unknown. A cellular stress response was captured by *USP44* and *WDR41*, genes linked to DNA damage control and nutrient sensing. Most notably, altered cell adhesion emerged as a recurring theme, spanning integrin-mediated cell-matrix attachment (*ITGA6*), epithelial junctional integrity (*PATJ*), and desmosomal adhesion (*DSC1*). ITGA6-containing integrins have been previously shown to be altered in localisation in SjD salivary glands [46,47]. Whether the circulating *ITGA6* alteration relates to these glandular changes remains unknown.

At the metabolomic level, 3-CMPFP, a furanoid fatty acid, had the largest coefficient in Score 2, and two further 3-CMPFP features and CMPF ranked among the top candidates. The structurally related CMPF is associated with metabolic and renal disease states [48] and has been linked to oxidative stress and glutathione depletion in experimental models [48,49]. Although exploratory, this finding suggests that redox homeostasis may differ between prevalent and incident lymphoma, in line with the recurrent glutathione/redox programmes identified here.

Several limitations should be acknowledged. Whole-blood microarray data capture a composite leukocyte signal, so some associations may partly reflect differences in cellular composition rather than cell-intrinsic transcriptional changes. The lymphoma groups were small, limiting statistical power, particularly for the metabolomic comparison of prevalent and incident lymphoma, and both the composite scores, evaluated only by LOOCV, and the lymphoma-associated pathway analyses require replication in independent cohorts. Differences in treatment exposure and renal function may also contribute to the circulating metabolomic profiles, while untargeted metabolomic annotation introduces uncertainty for some candidate features.

In conclusion, integrated transcriptomic and metabolomic profiling identified immune-metabolic programmes associated with SjD heterogeneity, including IFN/tryptophan, redox and innate inflammatory processes. Lymphoma was associated with an IL-18/IL-1β and oxidative stress transcriptional programme beyond clinical cluster membership, and prevalent lymphoma with an opposing IL-10/indole-related factor association in comparison to LSAHS. Shared transcriptomic features detected before diagnosis support further mechanistic investigation and independent prospective validation of blood-based biomarkers for lymphoma risk stratification.

## Ethics approval

This study was approved by the Ethics Committee of Bichat Hospital (Approval dossier N° 0711468). All patients gave written informed consent.

## Data availability statement

Anonymised data from the ASSESS cohort, along with the data dictionary, will be available upon publication. All requests for access must be directed to XM. Data transfer will be subject to an access protocol and will require authorisation from the scientific board of the ASSESS cohort. Data sharing will be contingent upon a data sharing agreement prior to any transfer, to ensure all users of the data adhere to the legal requirements of using personal data. Multi-omics analyses were performed with BiomiX (https://biomix-consortium.github.io/), whose source code is available at https://github.com/BiomiX-consortium/BiomiX2.5. Additional analysis scripts are available from the corresponding author upon reasonable request.

## Patient and public involvement

The project was presented and discussed with Pia Ducrey and Caroline Bühler, two patient research partners, during one of the regular consultation meetings, which are also used to identify research questions and topics of priority to the SjD patient community.

## Provenance and peer review

Not commissioned, externally peer reviewed.

## Declaration of generative AI use

Generative AI (Claude, Anthropic) was used to assist with language editing and proofreading of the manuscript text, and to assist with the development and modification of R scripts used for data analysis and figure generation. All AI-assisted output was reviewed, verified, and edited by the authors, who take full responsibility for the accuracy and integrity of the final manuscript and code.

## Contributorship (CRediT)

Conceptualization: B. Müller-Durovic, X. Mariette. Methodology: C. Iperi, S. Thing Teoh. Software: C. Iperi. Validation: C. Iperi, A. Kakale, S. Malkewitz, M. Sprecher. Formal analysis: C. Iperi, S. Thing Teoh. Investigation: C. Iperi. Resources: B. Müller-Durovic, X. Mariette, G. Nocturne, Y. Nguyen. Data curation: C. Iperi, S. Thing Teoh. Writing – original draft: C. Iperi. Writing – review & editing: all authors. Visualization: C. Iperi, A. Kakale, S. Malkewitz, M. Sprecher, B. Müller-Durovic, X. Mariette. Supervision: B. Müller-Durovic. Project administration: B. Müller-Durovic, X. Mariette, G. Nocturne, J.-E. Gottenberg. Funding acquisition: B. Müller-Durovic, X. Mariette.

## Acknowledgements

The authors thank all patients who participated in the ASSESS cohort and the investigators of the participating centres. We also thank our patient research partners: Pia Ducrey and Caroline Bühler, for ensuring that the study addresses patients’ needs.

