## Supplementary material, figures and tables for "Integrated transcriptomic and metabolomic profiling reveals distinct molecular programmes across clinical subgroups of Sjögren’s disease and signatures associated with lymphoma development": Supplementary_Figures.docx

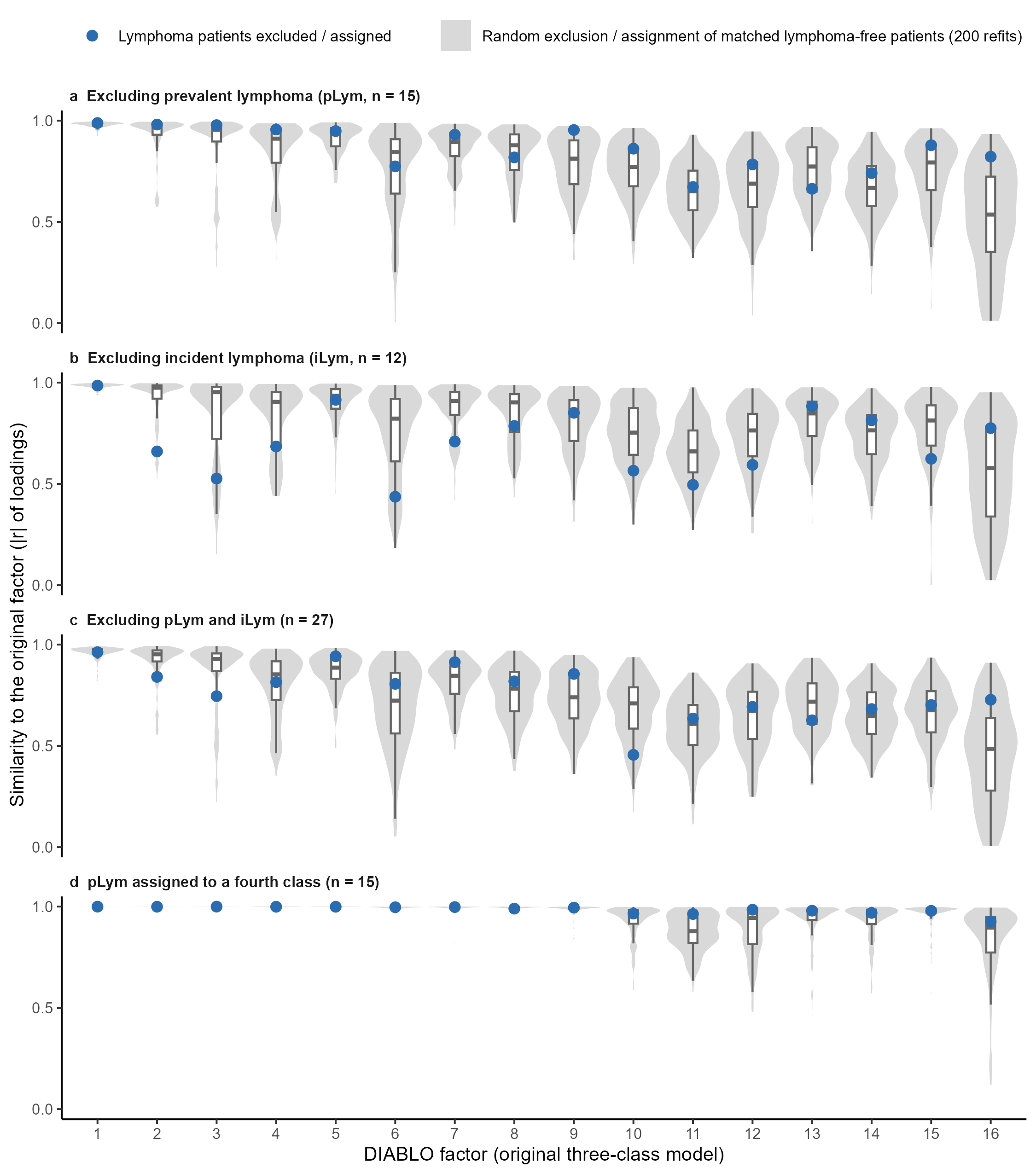

**Supplementary Figure 1. Influence of lymphoma patients on the DIABLO factor structure.** The three-class DIABLO model (BALS, HSA, LSAHS) was refitted with identical settings after excluding patients with (a) prevalent lymphoma (pLym, n=15), (b) incident lymphoma (iLym, n=12) or (c) both (n=27), and (d) after assigning pLym patients to a dedicated fourth class. Each refitted factor was matched one-to-one to the factors of the original three-class model (Hungarian algorithm), and similarity was measured as the absolute correlation (|r|) of factor loadings. Factor numbers refer to the original three-class model (Supplementary Table 2). Blue dots show the models in which lymphoma patients were excluded (a–c) or assigned to a fourth class (d). Grey violins and boxplots show the distribution obtained from 200 refits in which the same number of randomly selected lymphoma-free patients, with the same cluster composition, was excluded (a–c) or assigned to a fourth class (d). For all factors and all scenarios, observed similarity was within the range expected from random exclusion or assignment (all Benjamini–Hochberg-adjusted empirical p ≥ 0.35).
