## Supplementary material, figures and tables for "Integrated transcriptomic and metabolomic profiling reveals distinct molecular programmes across clinical subgroups of Sjögren’s disease and signatures associated with lymphoma development": Supplementary_methods_V1.9.docx

**Study cohort and participants**

This study was conducted within the ASSESS cohort, a prospective multicentre national cohort established in 2006 to identify predictors of systemic complications and lymphoma in Sjögren's disease during a 20-year follow-up [18,19]. Patients fulfilling the 2002 AECG criteria were recruited from 15 French tertiary centres between 2006 and 2009; the study was approved by the Ethics Committee of Bichat Hospital and all patients provided written informed consent. For the present analyses, 344 patients with available plasma metabolomic data were included; 306 also had matched whole-blood transcriptomic data and constituted the full multi-omics dataset. Patients were distributed across three previously defined clinical clusters (BALS, n=160; HSA, n=138; LSAHS, n=46) [18]. MOFA was applied to all 344 patients, exploiting its capacity to handle missing data blocks, whereas DIABLO was restricted to the 306 patients with complete multi-omics profiles. A subset of patients who had a lymphoma diagnosis at the time of cohort recruitment (pLym, n=17) or who developed lymphoma during follow-up (iLym, n=12) were analysed separately to characterise lymphoma-associated molecular signatures. Of these, 15 pLym and all 12 iLym had matched transcriptomic data and were included in DIABLO and transcriptome-based analyses.

**Transcriptomics data**

Whole-blood transcriptomic data were generated using Affymetrix GeneChip Clariom S Human microarrays as previously described by Trutschel et al. (GEO accession GSE140161) [24]. Transcriptomic profiling was performed at a single baseline time point. Normalised expression values were used as provided by the original publication, without additional transformation. For multi-omics integration analyses were selected the 5,000 most variable genes across all samples.

**Plasma metabolomics data acquisition and preprocessing**

Metabolites were extracted in deep 96-well plates. 20 µL plasma sample was aliquoted to each well, then mixed with 1000 µL of pre-chilled extraction solvent consisting of 40% acetonitrile, 40% methanol, 20% water. In total, 344 samples were extracted and distributed across four plates. The plates were centrifuged at 2200 x g, 4 C for 30 min, then 300 µL supernatant was transferred to a new plate. A further 10 µL was taken from each well, pooled together, re-centrifuged at 18000 x g, 4 C for 10 min, then aliquoted into several wells of the last plate (Plate 4) as QC samples. The plates were dried under nitrogen gas with an SC-200 Sample Concentrator (Cole-Parmer) then stored at -80 C until analysis.

For untargeted LC-HRMS analysis, the plates were thawed and analyzed sequentially, starting with Plate 4 containing the QC samples. Each well was reconstituted in 100 µL 80% water, 20 % acetonitrile. Prior to starting the entire run, QC samples were first re-pooled from individual wells and transferred to clear HPLC vials. One QC vial was run alongside each plate while the remaining vials were kept at -80 C until used.

Between the last two plates (Plate 2 and Plate 1, respectively), a dilution series of the QC sample (1, 2, 4, 8, 16x) was run analogously to the analytical samples, i.e. in full MS scan mode. This was later used for filtering peaks based on linear response of intensity vs dilution.

Metabolites were separated chromatographically on a Thermo Vanquish Horizon UHPLC system using a Waters ACQUITY HSS T3 column (2.1 × 150 mm) with a 1.8 µm particle size. A binary solvent system was employed, where Solvent A consisted of water with 0.1% formic acid and Solvent B consisted of acetonitrile with 0.1% formic acid. The column oven was maintained at 40 °C, and the autosampler tray was kept at 4 °C. Sample injection volume was 3 µL. Chromatography was performed with the following gradient program: t = 0 min, 0.250 mL/min flow rate, 0% B; t = 1 min, 0.300 mL/min flow rate, 0% B; t = 6 min, 0.300 mL/min flow rate, 20% B; t = 11 min, 0.300 mL/min flow rate, 100% B; t = 14 min, 0.450 mL/min flow rate, 100% B; t = 14.1 min, 0.250 mL/min flow rate, 0% B; t = 16 min , 0.250 mL/min flow rate. The chromatograph was coupled to a Thermo Q Exactive HF mass spectrometer via a heated electrospray ionization (HESI) source. The ionization parameters were set with spray voltages of +3.8 kV for positive mode and -3.5 kV for negative mode, an S-lens RF level of 50, and heated capillary temperature of 300 °C. The sheath gas flow rate was set to 30 and auxiliary gas to 10. Data acquisition of experimental samples was performed in full scan mode with polarity switching with mass range from m/z 70 to 1050, Orbitrap resolution of 60,000, automatic gain control (AGC) target of 3e6 and maximum injection time of 100 ms. Pooled QC samples were injected twice every 8 experimental samples for Data Dependent Acquisition (DDA), once each with positive and negative ionization modes respectively. The MS1 scan settings included mass range from m/z 70 to 1050, resolution of 60,000, AGC target of 1e6, and maximum injection time of 100 ms. MS2 acquisition settings included TopN of 5, intensity threshold of 1.6e5, isolation window of 1.0 m/z, dynamic exclusion window of 4.0 s, nominal collision energies (NCE) of 10, 20 and 40, dynamic m/z range based on precursor, resolution of 15,000, AGC target of 1e5, minimum AGC target of 8e3, and maximum injection time of 50 ms.

Raw data files were converted to mzML format using ProteoWizard msConvert [25]. Peak processing was performed separately for positive (full-scan analytical samples + positive DDA QC samples) and negative (full-scan analytical samples + negative DDA QC samples) modes using MZmine4 (https://mzio.io/mzmine). Briefly, processing steps included: Peak detection, chromatogram building and resolving, isotope grouping, peak alignment, gap filling, duplicate filtering, peak annotation by MS2 spectra [26]. MS2 spectral libraries used for peak annotation were from MassBank of North America (All LC-HRMS Orbitrap, downloaded on 31 July 2025) and MassBank EU (NIST format, downloaded on 17 June 2025). Exported peak areas were first filtered by linearity of intensity response vs relative concentration based on the QC dilution series injections. The filtered peaks were normalized by probabilistic quotient normalization (PQN) [27] then then log transformed to stabilise variance. Metabolite annotation was performed at both MS1 and MS2 levels using the BiomiX framework [28], which integrates CEU Mass Mediator [29] and TidyMass [30] for spectral matching against publicly available databases including the Human Metabolome Database (HMDB), KEGG, LipidMaps, METLIN, MassBank and the Mass Bank of North America (MoNA). Data Dependent Acquisition (DDA) MS2 fragmentation data were used to support annotation at the MS2 level where available. To account for technical sources of variation, batch effects attributable to plate, run order, and corticosteroid use (in the negative mode only) were removed using the removeBatchEffect function from the limma R package [31] prior to multi-omics integration.

**Unsupervised multi-omics factor analysis (MOFA)**

To characterise the major axes of biological variation within the ASSESS cohort in an unbiased manner, we applied Multi-Omics Factor Analysis (MOFA2, v1.14.0) [32] to the four matched data modalities: whole-blood transcriptomics (5,000 most variable genes), positive plasma metabolomics (1,413 peaks), negative plasma metabolomics (802 peaks), and clinical data (75 features). All features were variance-scaled prior to model fitting. The maximum number of latent factors was determined by an automatic stopping criterion, implemented within the BiomiX framework [28], whereby factor computation is halted when at least three consecutive factors explain less than 1% of variance in any single omics layer, ensuring that only factors with meaningful explanatory power are retained. To assess model stability and select the optimal number of latent factors, MOFA was run independently with five different random seeds. Model selection was performed using a composite score integrating four criteria: variance explained across omics layers (weight 25%), omic balance quantified as the coefficient of variation of variance explained across the three omics layers (weight 35%), sample clustering quality assessed by the mean silhouette score over k=2 to 8 clusters derived from the factor scores (weight 15%), and inter-factor redundancy quantified as the mean absolute pairwise correlation between factors (weight 25%, minimised). This weighting scheme was designed to prioritise models capturing balanced multi-omics information while minimising redundancy between factors. The composite score identified an optimal solution with 18 latent factors. Model stability was further validated by demonstrating consistent best silhouette scores across k=2 to 8 for all five seeds, and by computing the Adjusted Rand Index (ARI) between each seed model and a reference model (seed 42) across all tested factor numbers, confirming stable factor recovery up to 18 factors across independent model runs. Each factor represents a latent axis of co-variation shared across one or more omics layers and is defined by omic-specific feature weights (loadings), which quantify the contribution of each feature to that factor.

**Supervised multi-omics integration (DIABLO)**

To identify multi-omics signatures discriminating the three ASSESS clinical clusters, we applied the DIABLO framework (Data Integration Analysis for Biomarker discovery using Latent cOmponents) implemented in the mixOmics R package (v6.28.0) [33]. DIABLO simultaneously maximises the covariance between omics layers while optimising the separation between predefined clinical groups. The optimal number of latent components was selected by applying the same composite scoring criterion used for MOFA model selection, integrating variance explained, omic balance, sample clustering quality, and inter-factor redundancy, across candidate numbers of components. As DIABLO is deterministic, runs with different random seeds yield identical solutions; the robustness of the factors to patient composition was therefore assessed with the resampling procedure described below. Three types of DIABLO analyses were performed: (i) a primary analysis including patients assigned to the three clinical clusters with lymphoma patients embedded within their respective clusters; (ii) exclusion analyses in which patients with prevalent lymphoma (n=15), incident lymphoma (n=12) or both (n=27) were excluded, to assess the influence of lymphoma patients on the identified factors; and (iii) a four-group analysis in which the 15 patients with prevalent lymphoma were reassigned to a dedicated fourth cluster (pLym), to characterise lymphoma-specific molecular signatures in a supervised manner. As DIABLO cannot handle missing omics, all analyses were restricted to the 306 patients with complete matched transcriptomic and metabolomic profiles. The similarity of factors across DIABLO runs was quantified by computing the pairwise overlap between the top-contributing features of each factor, using the Jaccard index applied to the top-ranked loadings per omic per factor.

**Factor annotation**

Factors identified by both MOFA and DIABLO were annotated by systematic inspection of the top positive and negative loading features per omic per factor (scaled loadings > 0.5 or < −0.5, on a −1 to 1 scale). MOFA and DIABLO loadings were scaled to the range −1 to +1 by dividing by the maximum absolute loading across all views for each factor, so scores are comparable across features and views within a factor. For transcriptomic features, functional annotation and pathway enrichment analysis were performed using EnrichR [34], querying gene set libraries including GO Biological Process, KEGG, Reactome, and MSigDB Hallmarks. For metabolomic features, compound identity was assessed using the BiomiX annotation framework and, where necessary, cross-referenced against HMDB and the published literature. Factors dominated by technical signals (e.g. sex-linked variance, pharmaceutical compounds) were flagged and reported separately from biologically interpretable factors. Final biological annotations were assigned manually based on the convergent evidence from transcriptomic pathway enrichment and metabolomic compound identity.

**Four-group DIABLO and characterisation of prevalent lymphoma**

To characterise the molecular biology of lymphoma in the context of the biological axes identified by the three-group DIABLO analysis, the 15 patients with prevalent lymphoma and complete multi-omics data were reassigned to a dedicated fourth cluster (pLym) and DIABLO was rerun using the four-group design. As the resulting factor structure was highly concordant with the three-group solution, confirmed by pairwise feature overlap quantification, the four-group model allowed direct assessment of how pLym patients position relative to the three non-lymphoma clusters (BALS, HSA, LSAHS) across the same biological axes, without requiring reinterpretation of the factor identities.

**Influence of lymphoma patients on DIABLO factors**

To assess whether lymphoma patients shaped the DIABLO factor structure, the three-class model was refitted with identical settings after excluding patients with prevalent lymphoma, incident lymphoma or both, and after assigning patients with prevalent lymphoma to a dedicated fourth class. Factors of each refitted model were matched one-to-one to those of the original model using the Hungarian algorithm (clue R package) on the absolute correlation of block-normalised loadings. To obtain the variation expected from removing or relabelling any patient subset of the same size, each scenario was repeated 200 times with randomly selected lymphoma-free patients matched for cluster composition. For each factor, an empirical p-value was computed as the proportion of random refits with similarity lower than or equal to the observed one, and adjusted with the Benjamini-Hochberg procedure within each scenario (**Supplementary Figure 1**).

**Targeted analysis of lymphoma-associated programmes**

Programmes highlighted by DIABLO Factors 5, 7 and 8 were tested independently of DIABLO on the full whole-blood transcriptome (18,526 genes) rather than on the 5,000 most variable genes. Eight Gene Ontology gene sets were defined before testing and mapped to gene symbols with org.Hs.eg.db: IL-18/IL-1/inflammasome (GO:0035655, GO:0032621, GO:0044546, GO:0032611), glutathione metabolism (GO:0006749), and reactive oxygen and nitrogen species (GO:0072593, GO:0000302, GO:0006809). After the primary analysis, the glutathione gene sets originally used to annotate DIABLO Factors 7 (BioPlanet 2019: Glutathione conjugation; Glutathione metabolism) and 5 (Reactome 2024: Glutathione Synthesis and Recycling) were added from the Enrichr libraries, to identify the glutathione-related subprocess involved; false discovery rate for gene sets was computed across all 11 sets. For each patient, a pathway score was computed as the mean of the standardised expression of all genes in the set, and expressed in standard deviations of lymphoma-free patients. Scores were analysed with linear models including lymphoma status (none, prevalent, incident) and clinical cluster; the difference between prevalent and incident lymphoma was estimated from the same model. A combined comparison of all lymphoma patients (prevalent and incident) with lymphoma-free patients, adjusted for clinical cluster, was added as a secondary analysis after the separate comparisons, to increase statistical power. Factor 8 top-contributing genes (*IL18*, *IL18BP*, *IL1B*, *INPP4B*, *IER3*) were tested with limma, and 16 plasma features annotated as glutathione/thiol- or oxidative/nitrosative stress-related metabolites in Factors 5, 7 and 8 were tested with linear models, using the same design. False discovery rate was controlled with the Benjamini-Hochberg procedure separately for gene sets, genes and metabolites, within each comparison (**Supplementary Tables 4 and 5).**

**Unbiased discriminative screening, composite score development and cross-validation**

To identify molecular features associated with lymphoma risk in an unbiased manner independent of the DIABLO factor structure, twelve incidence lymphoma patients (iLym) were merged in a separate group. A Kruskal-Wallis test was applied to every available variable across the complete transcriptomic and metabolomic matrices across all five groups (BALS, HSA, LSAHS, pLym, iLym), and variables were ranked by their global p-value. The top 50 discriminative features were inspected for each omic layer separately to guide variable selection for composite score development. Two composite molecular scores were developed to discriminate lymphoma-risk patients from non-lymphoma patients and, secondarily, to distinguish incident from prevalent lymphoma. Score 1 was built using 35 transcriptomic features selected from the genome-wide Kruskal-Wallis screening as the top discriminators of lymphoma (pLym + iLym) versus no lymphoma (BALS, HSA, LSAHS). Score 2 was built using 15 plasma metabolomic features (11 positive mode, 4 negative mode) selected as the top discriminators of pLym versus iLym.

Each score was modelled using L1-penalised logistic regression (LASSO) implemented in the glmnet R package (v4.1-8), with inverse class-frequency sample weighting to account for class imbalance (279 non-lymphoma vs 27 lymphoma-risk patients for Score 1). Model performance was evaluated using Leave-One-Out Cross-Validation (LOOCV): at each iteration, the model was trained on N−1 samples with feature scaling (z-score normalisation) computed exclusively on the training partition and applied to the left-out sample to generate a predicted probability. The predicted probabilities from all LOOCV iterations were aggregated to construct a single pooled ROC curve, from which the Area Under the Curve (AUC) was derived. The 95% confidence interval for the AUC was estimated by bootstrap resampling (1,000 iterations) using the ci.auc function from the pROC R package. The optimal classification threshold was determined by maximising the Youden Index (sensitivity + specificity − 1). To assess whether model performance exceeded chance, a permutation test was performed: the full LOOCV procedure was repeated 1,000 times with randomly permuted class labels, generating a null distribution of AUC values. The permutation p-value was defined as the proportion of null AUC values equal to or exceeding the observed AUC.

**Statistical analyses and software**

All analyses were performed in R (v4.4.3). Key packages included MOFA2 (v1.14.0), mixOmics (v6.28.0), glmnet (v4.1-8), pROC, limma (v3.50.3), clue (v0.3-64), org.Hs.eg.db (v3.14.0), ggplot2, and ggpubr. Kruskal-Wallis tests and post-hoc Dunn tests with Benjamini-Hochberg correction were used for multi-group comparisons. Wilcoxon rank-sum tests were used for pairwise comparisons. A two-sided significance threshold of p < 0.05 was applied throughout unless otherwise stated.
